# Bone innervation and vascularization regulated by osteoclasts contribute to refractive pain-related behavior in the collagen antibody-induced arthritis model

**DOI:** 10.1101/2021.04.19.440384

**Authors:** Resti Rudjito, Nilesh M Agalave, Alex Bersellini Farinotti, Azar Baharpoor, Arisai Martinez Martinez, Enriqueta Muñoz Islas, Preety Panwar, Dieter Brömme, Julie Barbier, Fabien Marchand, Patrick Mehlen, Thomas Levin Andersen, Juan Miguel Jimenez Andrade, Camilla I. Svensson

## Abstract

**Objective:** Rheumatoid arthritis is often characterized by eroded joints and chronic pain that outlasts disease activity. Whilst several reports show strong associations between bone resorption and nociception, the underlying mechanisms remain to be unraveled. Here, we used the collagen antibody-induced arthritis (CAIA) model to examine the contribution of osteoclasts in pain regulation. The antinociceptive effects of osteoclasts inhibitors and their mechanisms of actions involving bone vascularization and innervation were also explored.

**Methods:** BALB/c female mice were subjected to CAIA by intravenous injection of a collagen type-II antibody cocktail, followed by intraperitoneal injection of lipopolysaccharide. Degree of arthritis, bone resorption, mechanical hypersensitivity, vascularization and innervation in the ankle joint were assessed. Animals were treated with osteoclast inhibitors, zoledronate and cathepsin K inhibitor (T06), and netrin-1 neutralizing antibody. Potential pronociceptive factors were examined in primary osteoclast cultures.

**Results:** CAIA induced local bone loss in the calcaneus with ongoing increased osteoclast activity during the inflammatory phase of the model, but not after inflammation has resolved. Mechanical hypersensitivity was reversed by zoledronate in late but not inflammatory phase CAIA. This effect was coupled to the ability of osteoclasts to modulate bone vascularization and innervation, which was inhibited by osteoclast inhibitors. CAIA-induced hypersensitivity in the late phase was also reversed by anti-netrin-1 antibody.

**Conclusion:** Osteoclasts induce pain-like behavior in the CAIA model independent of inflammation via effects on bone vascularization and innervation.

**Key messages:** What is already known about this subject?

- Pain and residual signs of erosive lesions are frequently present in rheumatoid arthritis (RA) patients with good disease control
- Osteoclasts can induce nociceptive signaling but the exact mechanism with respect to RA-induced pain is not clear

What does this study add?

- The pronociceptive actions of osteoclasts extend beyond flares of joint inflammation and erosive activity by increasing bone innervation, bone vascularization and netrin-1 release
- Osteoclast inhibitors and neutralizing netrin-1 antibodies reverse refractive pain-related behaviors in the collagen antibody-induced arthritis model

How might this impact on clinical practice or future developments?

- This study provides insights to the potential of osteoclast inhibition as a therapeutic strategy for persistent pain in RA

## INTRODUCTION

Rheumatoid arthritis (RA) is a chronic autoimmune disease, typically presented as symmetrical pathology in small synovial joints and accompanied by symptoms such as joint pain (arthralgia), swelling and fatigue that may have debilitating effects on the functional ability and quality of life [1, 2]. The experience of pain for individuals with RA is highly varied and do not completely covary with inflammation. Patients can present arthralgia before showing visual signs of inflammation [1, 3], and many report pain even after their disease activity is well controlled via pharmacological interventions [4, 5].

While the inflamed synovium has traditionally been considered as the sole elicitor of pain in RA, the frequent misalignment between inflammation and pain has encouraged researchers to search for additional mechanisms that drive pain in arthritis. The notion that structural changes in bone such as focal and subchondral bone loss are associated with pain has underbuilt the hypothesis that osteoclasts play a critical role in the regulation of processes leading to sustained pain. Indeed, preclinical and clinical studies have demonstrated the benefits of osteoclast inhibitors, including bisphosphonates and calcitonin, in alleviating hyperalgesia in multiple bone disorders [6–8]. Osteoclasts degrade the mineralized bone matrix by secreting protons and acidify the extracellular environment [9], which is a condition known to be algogenic for nociceptive neurons [10, 11]. Interestingly, reports also show that specific osteoclast inhibitors can attenuate pain-like behavior in models that do not exhibit bone loss [12–14]. This suggests that the pronociceptive properties of osteoclasts and the actions of osteoclast inhibitors are not necessarily coupled to resorption, but could be mediated via non-resorbing actions of osteoclasts, potentially by releasing inflammatory, angiogenic and neurotrophic factors such as interleukin (IL)-8, platelet-derived growth factor (PDGF-BB), semaphorin (Sema)4D and netrin-1 [15–19]. Hence, there are several ways osteoclasts may contribute to pain and the mechanism by which these cells regulate nociception in RA warrants further investigation.

For this purpose, we used the collagen antibody-induced arthritis (CAIA) model in which mechanical hypersensitivity was previously observed prior to, during, and following the antibody-induced flare of joint inflammation [20–25]. Interestingly, signs of bone loss remain several weeks after resolution of inflammation [20], raising the question whether bone erosion contribute to the persistency of hypersensitivity in this model. Thus, the aims of this study were to i) characterize changes in bone metabolism in the CAIA model, ii) examine the effect of osteoclast inhibition in CAIA-induced hypersensitivity and iii) explore through which mechanisms these inhibitors exert their antinociceptive effects.

## MATERIALS AND METHODS

Additional detailed information is described in the supplementary file.

### Animals experiments

BALB/c female mice (10-12 weeks, 20-25 g) were purchased from Charles River and Janvier laboratories and housed in climate-controlled environment (5 mice per cage) with 12 h light/dark cycle and *ad libitum* access to food and water. All experiments conformed to the ARRIVE guidelines and protocols approved by local ethical committees in Sweden (Stockholm Norra Djurförsöksetiska nämnd) and France (CEMEA-Auvergne).

CAIA was induced by intravenous (i.v.) injection of a collagen type II (CII) arthritogenic antibodies (1.5 mg/mouse, Chondrex) on day 0 followed by intraperitoneal (i.p.) injection of lipopolysaccharide (LPS, 25 μg/mouse, Chondrex) on day 3 or 5 to synchronize inflammation. LPS injection on day 3 gives higher arthritis incidence and therefore the protocol was refined. The control group received saline i.v. on day 0 and i.p. on day 3 or 5. Arthritis scoring of the fore and hind paws were performed as previously described [20]. A score of 1 point was given for each inflamed toe or knuckle and a score of maximum 5 points was given to inflamed wrist or ankle. The effect of subcutaneous (s.c.) injection of zoledronate (100 μg/kg, every 3 days), oral administration of the cathepsin K inhibitor tanshinone IIA sulfonic sodium (T06, 40 mg/kg, daily) or i.p. injection of anti-Netrin-1 antibody (Net1-mAb, NP137, 10 mg/kg, every 2 days) were investigated in the CAIA model.

### Mechanical hypersensitivity

Mechanical hypersensitivity was assessed by application of calibrated von Frey filaments (Marstock) as previously described [20]. A 50% paw withdrawal threshold was calculated using the up-down method [26]. The experimenter was blinded to the treatment groups during the behavioral tests.

### Tissue analysis

Micro-computed tomography (micro-CT) imaging, scintigraphic imaging, immunoassay, immunohistochemistry (IHC), *in situ* hybridization (ISH) and qPCR analyses on serum, ankle joints and dorsal root ganglia (DRGs) are described in supplementary information.

### Osteoclast cultures

Bone marrow cells obtained from mouse femur and tibia were differentiated into osteoclasts with M-CSF and RANKL (both R&D) stimulation. Cells were cultured with either 600 nM zoledronate (Sigma-Aldrich), 1 μM T06 or vehicle (PBS). PDGF-BB and Sema4D were quantified in culture supernatant. At the end of the experiment, cells were either fixed with 4% paraformaldehyde for quantification of TRAP^+^ osteoclasts or subjected to protein extraction for measurement of netrin-1 levels using Western Blot.

### Primary neuronal cultures

DRG neuronal cultures were prepared using a modified version of previously described protocol [27]. Briefly, mouse DRGs (C1-L6) were dissected and enzymatically digested with papain and collagenase I/dispase II. Isolated cells were then seeded on glass-bottomed dishes pre-coated with poly-D-lysine (Corning) and laminin (Sigma-Aldrich) and kept in a humidified 37°C incubator with 5% CO_2_ for 24 h. After 24 h, calcium imaging was performed to determine intracellular calcium cell responses to direct application of mouse recombinant netrin-1 (R&D).

### Statistical analysis

For comparing changes over time, two-way analysis of variance was used followed by Bonferroni post hoc test. Mean differences between three groups or more were compared using one-way analysis of variance followed by Bonferroni or Dunnett post hoc test, while differences in two groups were analyzed by student’s *t*-test. All tests were performed using GraphPad Prism software. p values < 0.05 were considered statistically significant.

## RESULTS

### CAIA induced bone erosion

MicroCT analyses showed robust trabecular bone loss in the calcaneus, detected by decreased bone mineral density (BMD), bone volume per tissue volume (BV/TV) and trabecular number (Tb.N), as well as increased trabecular separation (Tb.Sp), in mice subjected to CAIA day 17-20 and 54-56 compared to saline control (Figure 1A-D, Table S1). Changes in trabecular thickness (Tb.Th) were, however, not detected in all groups for both phases (Figure 1C, D). No significant correlations were observed between arthritis scores and severity of erosive lesions in CAIA-subjected animals (Figure S1A, B).

**Figure 1.**
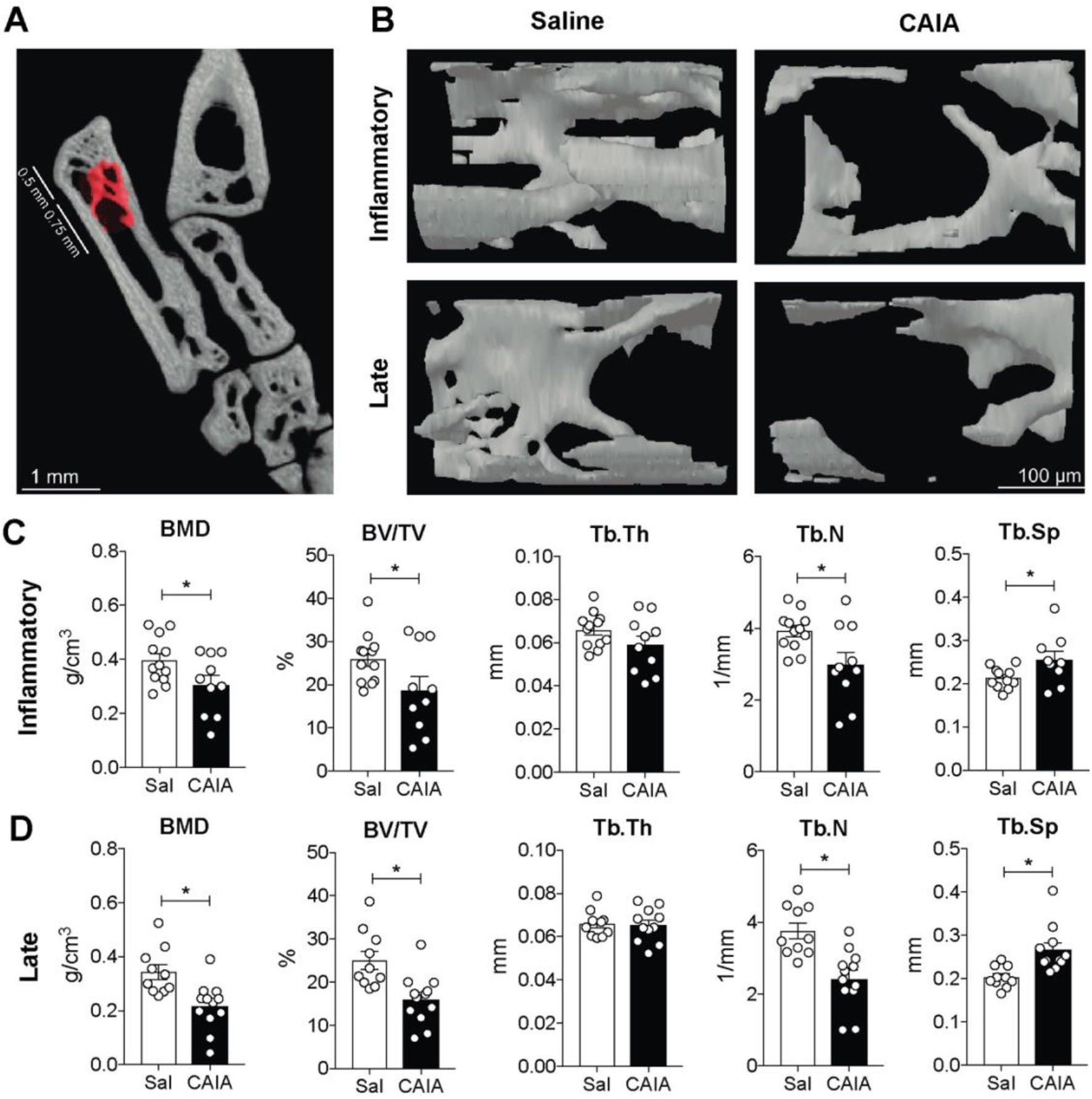
Collagen antibody-induced arthritis (CAIA) promotes localized bone loss in the calcaneus. Ankle joints were harvested from the inflammatory (day 17-20) and late (day 54-56) phase of the CAIA model. Representative micro-CT images of quantification area in the calcaneus **(A)** and images of calcaneus from saline and CAIA mice in inflammatory and late phase **(B)**. Bar graphs show quantitative evaluation of trabecular bone mineral density (BMD), bone volume per tissue volume (BV/TV), trabecular number (Tb.N), trabecular thickness (Tb.Th) and trabecular separation (Tb.Sp) in calcaneus of saline and CAIA mice in inflammatory **(C)** and late **(D)** phase. Data are presented as mean ± SEM, n=9-12 mice/group, *p<0.05, **p<0.01, ***p<0.001, Sal: saline control.

### Osteoclast activity was enhanced in the inflammatory phase of the CAIA model

We next determined if osteoclast activity in the CAIA model is continuously elevated or if there is residual bone loss detected in the late phase. First, evaluating *Acp5* and *Ctsk* mRNA levels in whole ankle joints showed no alterations in both phases of the CAIA model compared to control groups (Figure 1A). Second, we quantified osteoclast numbers specifically in the calcaneus as changes in osteoclast activity may be local and not detectable in bulk mRNA from whole joint extracts. A significant increase in *Acp5*^+^ and CTSK+ osteoclast surface per bone surface (Oc.S/B.S) was observed in the calcaneus of CAIA mice compared to the control in the inflammatory phase, but not the late phase (Figure 2B, C). Next, we examined changes in bone remodeling. Serum levels of the bone resorption marker CTX-I were elevated in the inflammatory phase, but the levels of the bone formation marker PINP were not altered in either phases (Figure 2D). Furthermore, ^99m^Tc-HMDP scintigraphic imaging of the ankle joint showed elevated levels of ^99m^Tc-HMDP uptake in CAIA mice only on day 17 but not at later timepoints (Figure 2E), thus suggesting a higher bone turnover during inflammation. Lastly, we evaluated expression of genes associated with acidification as this mechanism has been attributed to osteoclast-induced nociception [10, 11]. We found upregulation of *Tcirg1* and *Clcn7* mRNA levels in ankle joints from inflammatory but not late phase CAIA mice (Figure 2F), but the levels of *Asic3* and *Trpv1* in DRGs remained unchanged at all timepoints (Figure 2F).

**Figure 2.**
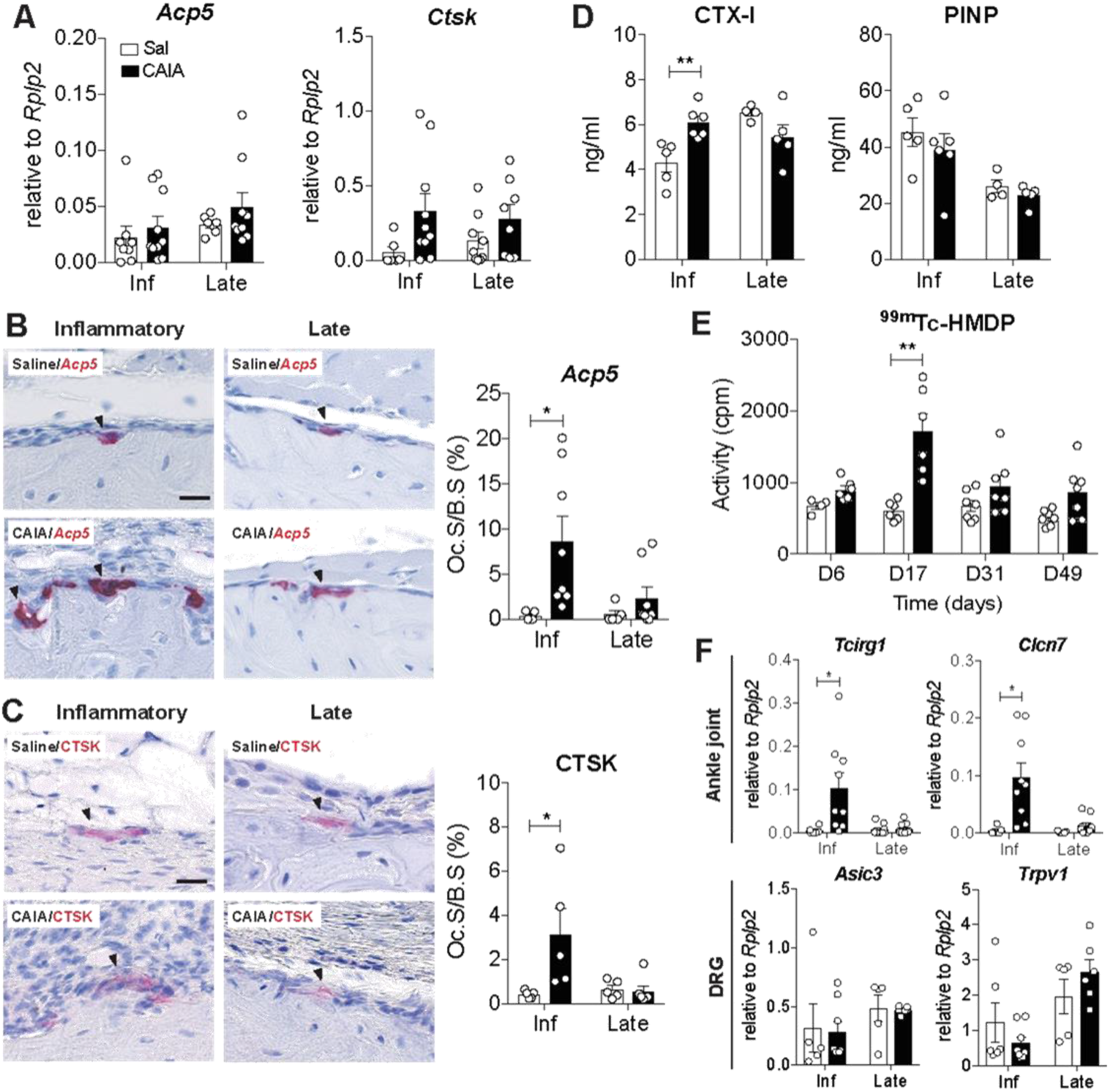
Enhanced osteoclast activity occurs in the inflammatory but not late phase of the collagen antibody-induced arthritis (CAIA) model. Ankle joint and serum samples were collected from the inflammatory (day 17-20) and late (day 54-56) phases. Bar graphs show mRNA levels for common osteoclast markers *Acp5* and *Ctsk* in the ankle joint **(A)**. Representative images and quantification of tartrate-resistant acid phosphatase (*Acp5*) *in situ* hybridization **(B)** and cathepsin K (CTSK) immunohistochemical staining **(C)** in the calcaneus, expressed as osteoclast surface per bone surface (Oc.S/BS). Arrowheads indicate positively-stained cells, scale bar: 25 μm. Bar graphs represent C-terminal telopeptide of collagen type I (CTX-I, bone erosion marker) and N-terminal propeptide of collagen type I (PINP, bone formation marker) **(D)** levels in serum. Bar graph represents ^99m^TcHMDP uptake on day 6, 17, 31, 49 of the CAIA model, which assesses for bone remodeling activity **(E)**. Bar graphs show mRNA levels for acidification-associated genes *Tcirg1* and *Clcn7* (upper panel) and acid sensing channels *Asic3* and *Trpv1* (lower panel) in the ankle joint and dorsal root ganglion (DRG), respectively (**F**). Data are presented as mean ± SEM, n=4-11 mice/group, *p<0.05, **p<0.01. Sal: saline control, Inf: inflammatory phase.

### Osteoclast inhibitors reversed CAIA-induced hypersensitivity in the late phase

To decipher the link between osteoclasts and pain, two different osteoclast inhibitors were assessed in the CAIA model. Systemic continuous zoledronate administration (100 μg/kg) on days 6-56 did not affect arthritis scores, but led to a progressive reversal of mechanical hypersensitivity in the late phase, which was significant from day 40 onwards (Figure 3A, B). To verify that this antinociceptive effect was restricted to the late phase, zoledronate was administered from days 6-19 or days 43-56. In agreement, paw withdrawal thresholds were normalized only with treatment in the late phase (Figure 3C, D). Administration of T06 (40 mg/kg) during the inflammatory phase (days 6-20) or late phase (days 43-59) evoked similar effects as zoledronate. The degree of arthritis and mechanical hypersensitivity were not altered when T06 was delivered during inflammation (Figure 3E), but a significant reversal of the mechanical hypersensitivity was observed in the late phase (Figure 3F). Treatments with zoledronate (days 6-56) or T06 (days 43-59) resulted in an increase in BMD, BV/TV and Tb.N, and decrease in Tb.Sp in the calcaneus (Figure 4A-E) compared to vehicle-treated CAIA mice.

**Figure 3.**
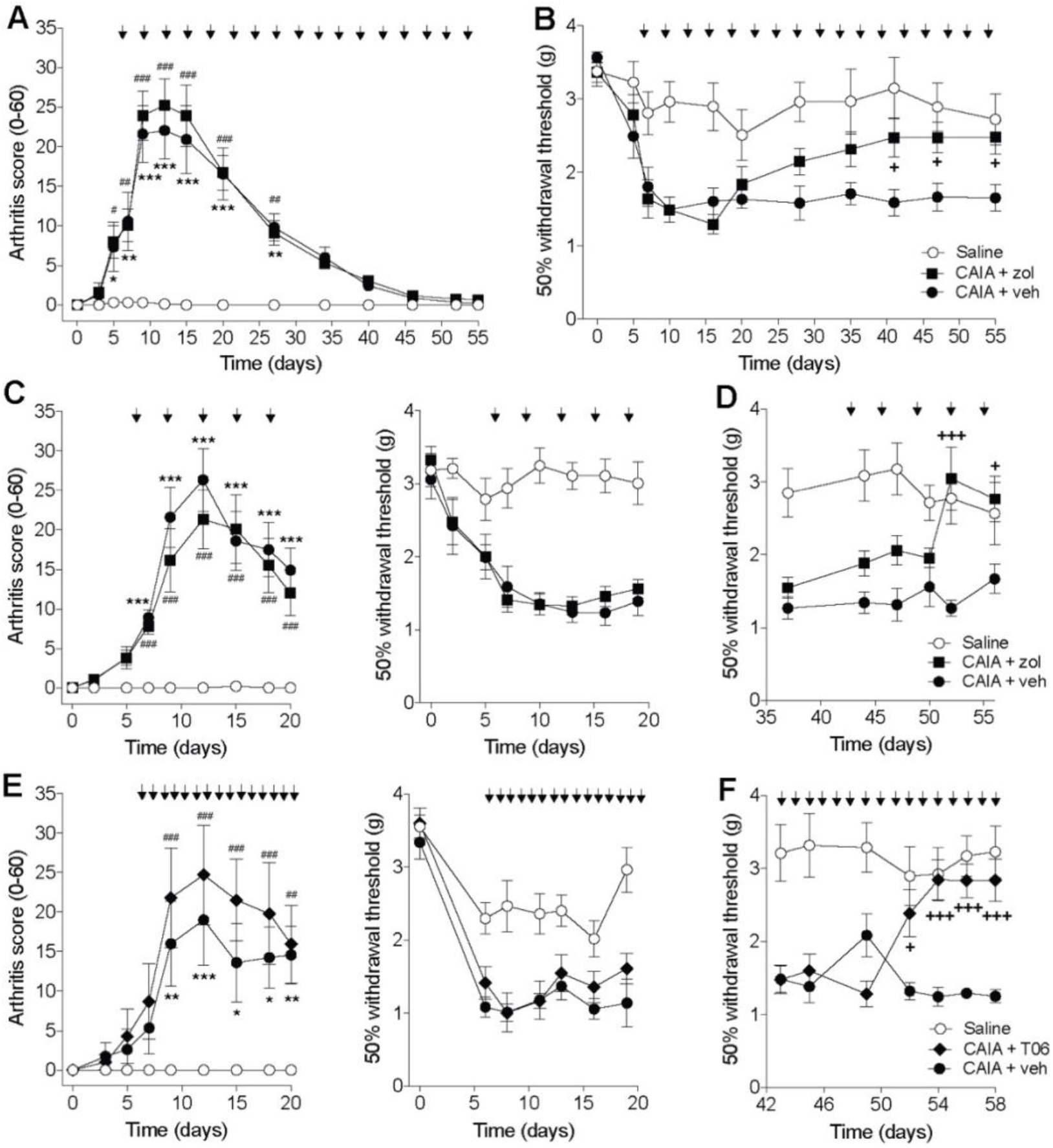
The osteoclast inhibitors zoledronate and T06 reverse mechanical hypersensitivity only in the late phase of the collagen antibody-induced arthritis (CAIA model) without diminishing joint inflammation. Line graphs represent the degree of arthritis assessed by visual inspection of the hind ankle joints (arthritis score 0-60) **(A)** and withdrawal thresholds assessed by von Frey filaments **(B)** following chronic administration of zoledronate s.c. (100 mg/kg) once every third day between day 6-56. Line graphs illustrate arthritis scores and withdrawal threshold to mechanical stimulation subsequent to acute zoledronate treatment in the inflammatory phase (day 6-19) (**C**) and late phase (day 43-56) **(D)**. Line graphs depict arthritis severity by visual inspection of the hind ankle joints and mechanical withdrawal thresholds following T06 (40 mg/kg) administration by daily oral gavage on day 6-20 **(E)** and day 43-59 **(F)**. Data are presented as mean ± SEM, n=5-11 mice/group, ^*,#,+^p<0.05, ^**,##^p<0.01, ^***,###,+++^p<0.001 where *represents saline vs. CAIA + vehicle, #represents saline vs. CAIA + osteoclast inhibitor and ^+^represents CAIA + osteoclast inhibitor vs. CAIA + vehicle. Arrows indicate drug administration.

**Figure 4.**
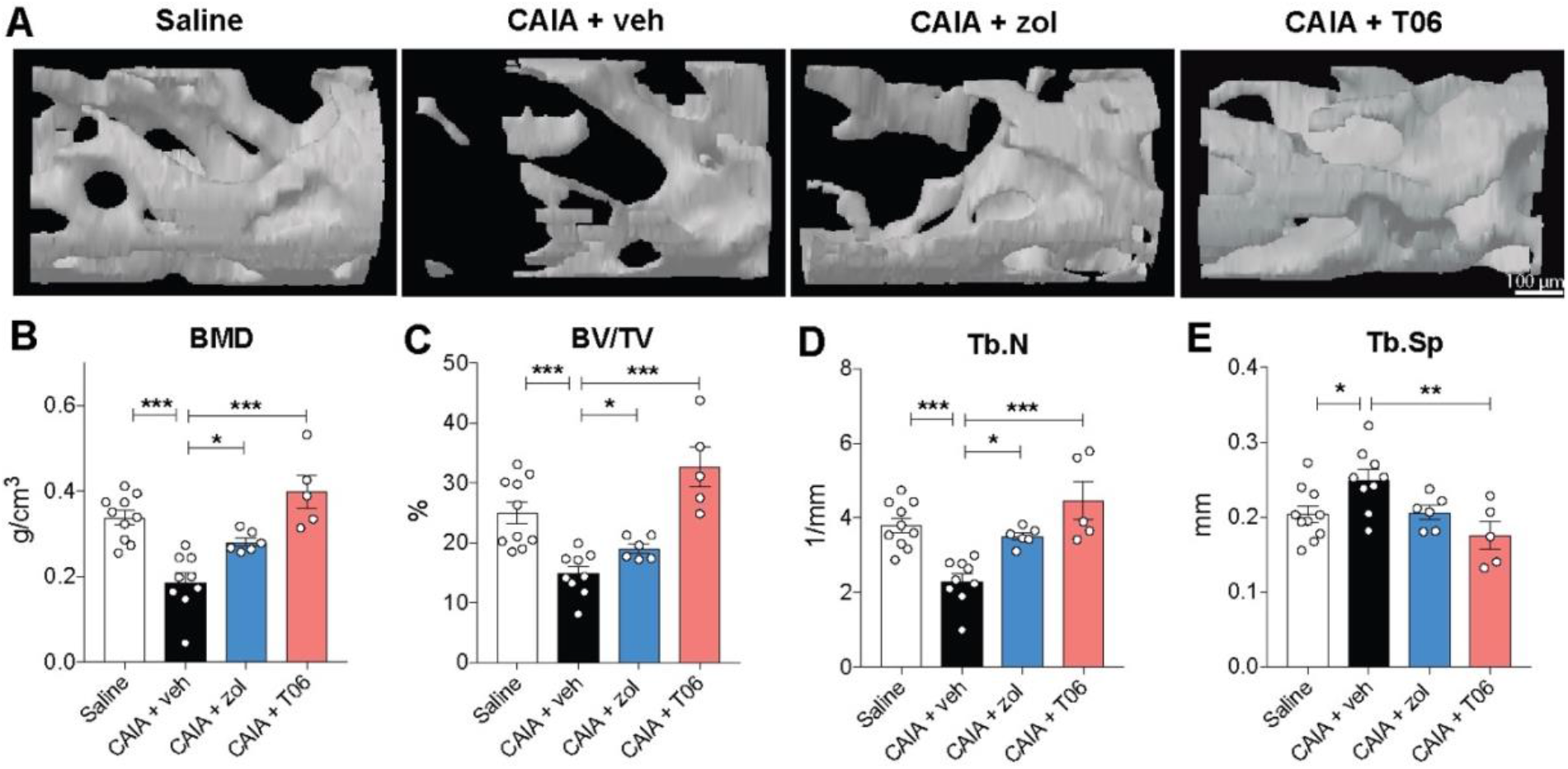
Zoledronate and T06 reduced CAIA-induced bone loss in the late phase. Ankle joints were harvested from the late phase (day 56-59) of the CAIA model. Representative micro-CT images of calcaneus from saline control, CAIA + vehicle (veh), CAIA + zoledronate (zol) and CAIA + T06 groups **(A)**. Scatter graphs show quantitative evaluation of trabecular bone mineral density (BMD) **(B)**, bone volume per tissue volume (BV/TV) **(C)**, trabecular number (Tb.N) **(D)** and trabecular separation (Tb.Sp) **(E)** in the calcaneus. Data are presented as mean ± SEM, n=5-10 mice/group, *p<0.05, **p<0.01, ***p<0.001.

### Osteoclast inhibitors altered CAIA-induced vascularization and innervation

In order to identify alternative mechanisms by which osteoclasts can mediate nociception, we analyzed mRNA levels for the endothelial marker *Pecam1* and the axon guidance molecules *Sema4d* and *Ntnl1* in whole ankle joint extracts. *Pecam1* expression was upregulated in both inflammatory and late phase CAIA, but was normalized to control levels by either zoledronate or T06 in the late phase (Figure 5A). In contrast, *Ntn1 and Sema4d* mRNA levels were only increased in the late phase and decreased by either zoledronate or T06 administration, though the reduction of *Sema4d* expression did not reach statistical significance following zoledronate treatment (Figure 5A). Next, IHC was performed to examine vascularization and innervation in the periosteum of the calcaneus. Vascular and nerve densities were estimated by quantifying the number of CD31^+^ vessels or PGP9.5^+^ nerve profiles within 100 μm from the surface of calcaneus (Figure 5B, D). CAIA increased vascular densities in both inflammatory and late phases compared to the control group, which was significantly reduced by both either zoledronate and T06 only in the late phase (Figure 5C). An increase in nerve profile density was also observed in both phases of the CAIA model compared to the control groups, which was significantly decreased by the two osteoclast inhibitors in the late phase (Figure 5E).

**Figure 5.**
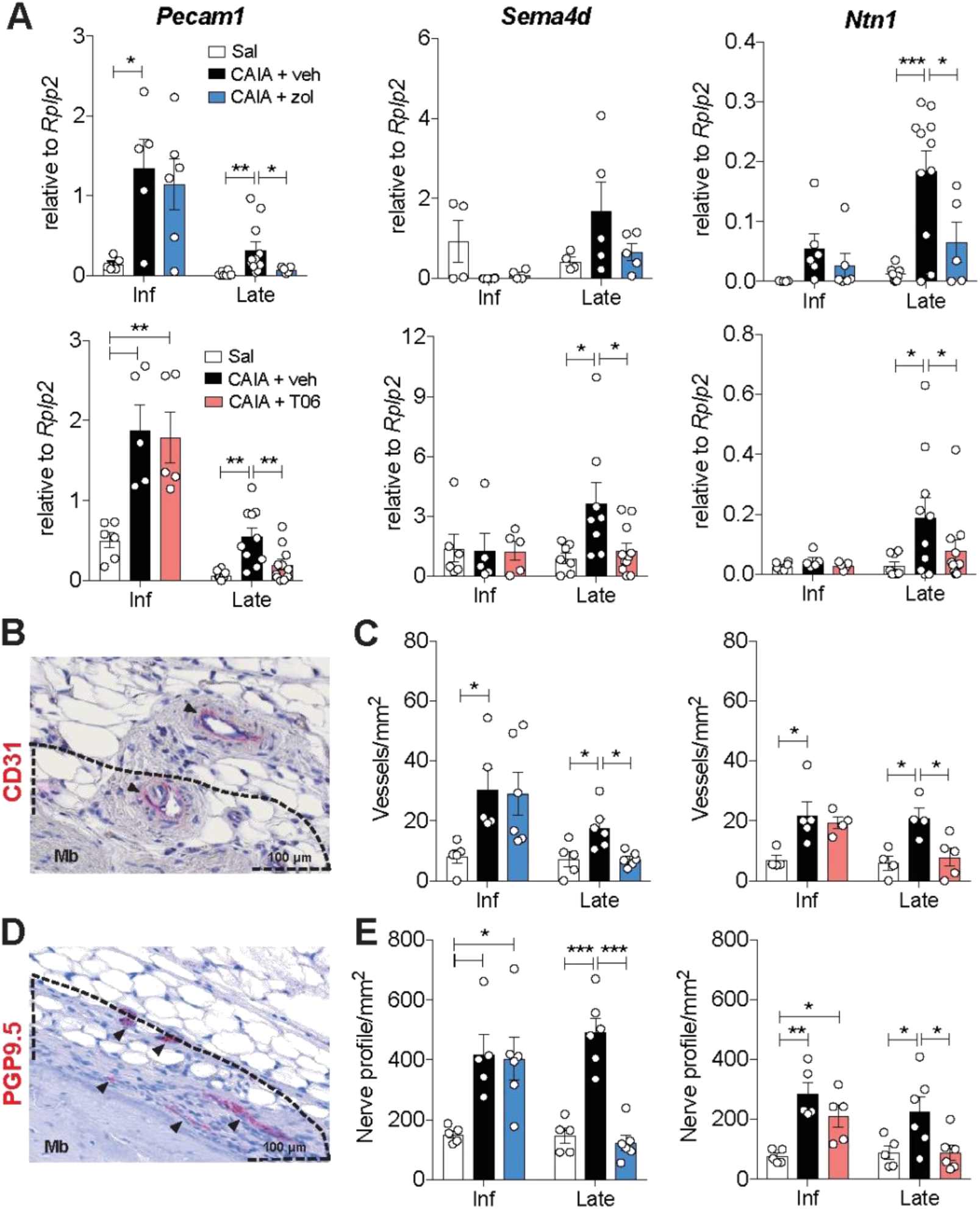
Zoledronate and T06 decreased collagen antibody-induced arthritis (CAIA)-induced neovascularization and nerve fiber sprouting in the calcaneus only in the late phase. Ankle joints were harvested from the inflammatory (day 17-20) and late (day 54-59) phase of the CAIA model. Bar graphs represent mRNA levels for *Pecam1, Sema4d*, and *Ntn1* in inflammatory and late phase CAIA joints treated with zoledronate or T06 **(A).** Ankle joint sections were stained for CD31 (endothelial marker) or PGP9.5 (neuronal marker) and vascular and nerve profile densities were evaluated in the calcaneus. Representative immunohistochemical images of quantification area for CD31 **(B)** and PGP9.5 **(C)**, which is < 100 μm from the surface of the calcaneus. Arrowheads indicate positive staining. Bar graphs represent the vascular **(D)** and nerve profile densities **(E)** in saline control (sal), CAIA + vehicle (veh), CAIA + zoledronate (zol) and CAIA + T06 mice in both inflammatory and late phase. Data are presented as mean ± SEM, n=4-11 mice/group, *p<0.05, **p<0.01, ***p<0.001. Inf: inflammatory phase, Mb: mineralized bone.

### Osteoclasts produced angiogenic and neurogenic factors

To elucidate if osteoclasts produce factors that may modulate vascularization and innervation, primary osteoclasts were cultured with either vehicle (PBS), zoledronate (600 nM) or T06 (1 μM). Differentiation with both M-CSF and RANKL resulted in TRAP^+^ multinucleated cells on day 6, which are common characteristics of osteoclasts, as opposed to cells cultured with M-CSF only that did not generate cells with this morphology (Figure 6A, B). TRAP^+^ osteoclast number was significantly decreased following zoledronate but not T06 treatment (Figure 6A, B). Next, we examined if osteoclast inhibition affects the production of PDGF-BB, Sema4D and netrin-1. Osteoclast differentiation led to significant increase in PGDF-BB, Sema4D and netrin-1 protein levels (Figure 6C-E). PDGF-BB levels were significantly reduced by both zoledronate and T06 (Figure 6C), while the reduction of Sema4D and netrin-1 protein levels only reached statistical significance in the presence of zoledronate and not T06, compared to vehicle-treated cells (Figure 6B-D).

**Figure 6.**
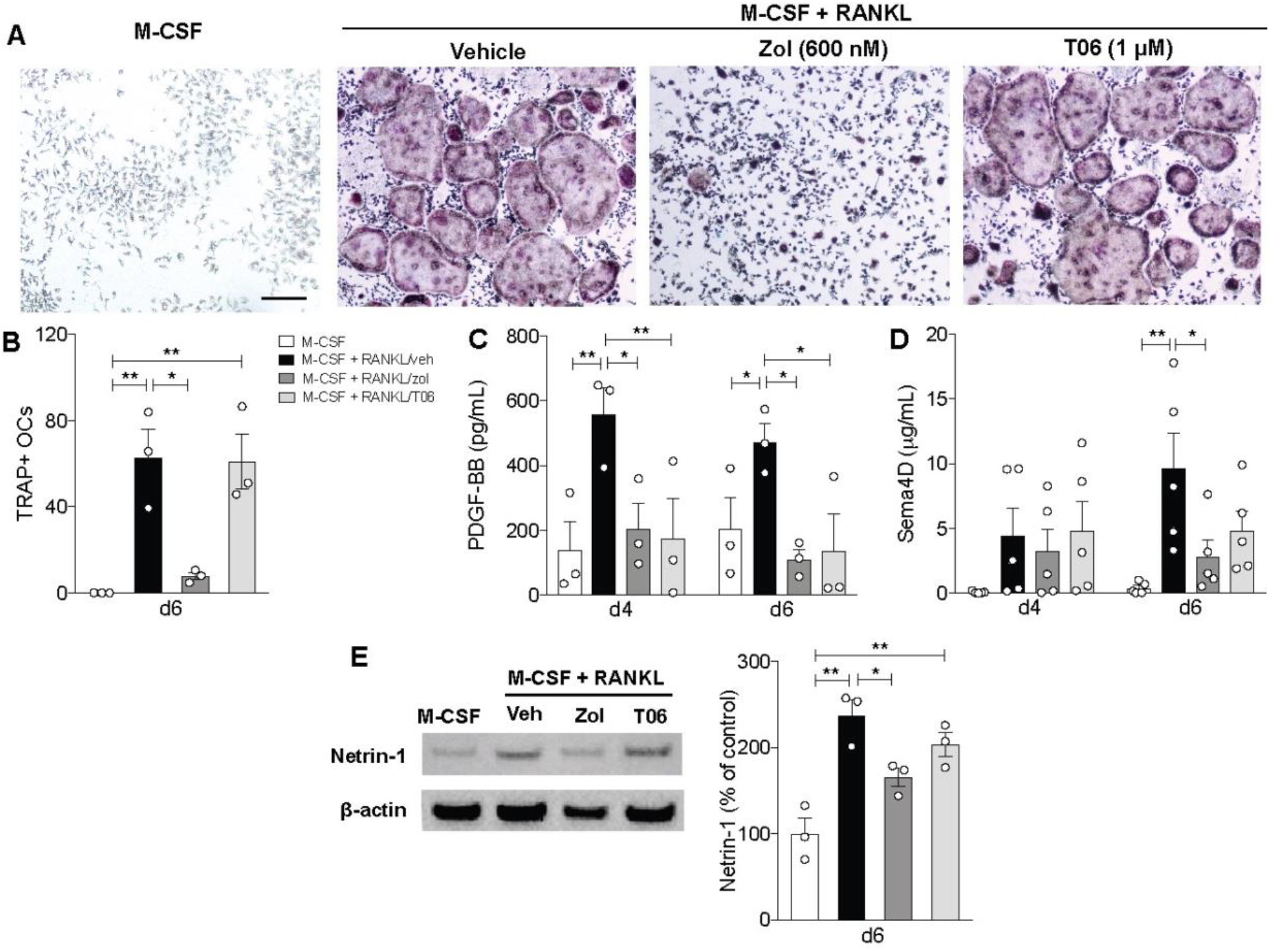
Osteoclasts produce angiogenic and neurogenic factors which are inhibited by either zoledronate or T06. Representative images of TRAP-stained cells **(A)** and quantification of TRAP^+^ multinucleated cells (>3 nuclei) **(B)** following treatments with macrophage colony stimulating factor (M-CSF) only or a combination of M-CSF and receptor activator RANKL supplemented with either vehicle (veh), 600 nM zoledronate (zol) or 1 μM T06, scale bar: 100 μm. Bar graphs represent the level of PDGF-BB **(C)** and Sema4D **(D)** measured in osteoclast culture supernatant collected on day 4 and 6. Western Blot images and quantification of netrin-1 on day 6 osteoclast culture **(E)**. Data are presented as mean ± SEM, n=3-5 independent experiments, *p<0.05, **p<0.01.

### Netrin-1 regulated CAIA-induced nociception

Recent work suggests that netrin-1 plays a crucial role in bone pain [15, 28, 29] and therefore we examined the contribution of this factor to CAIA-induced pain-like behavior. Systemic administration of the anti-netrin-1 antibody (NP137, 10 mg/kg) did not affect arthritis score or mechanical hypersensitivity in the inflammatory phase (Figure 7A, B). However, we observed a reversal of CAIA-induced hypersensitivity in the late phase with this drug starting from day 49 onwards (Figure 7C). To further confirm the pronociceptive property of netrin-1, *in vitro* calcium imaging experiments with primary DRG neurons were performed. Netrin-1 stimulation increased the percentage of DRG neurons that showed elevated intracellular Ca^2+^ signal in a dose-dependent fashion (Figure 7D).

**Figure 7.**
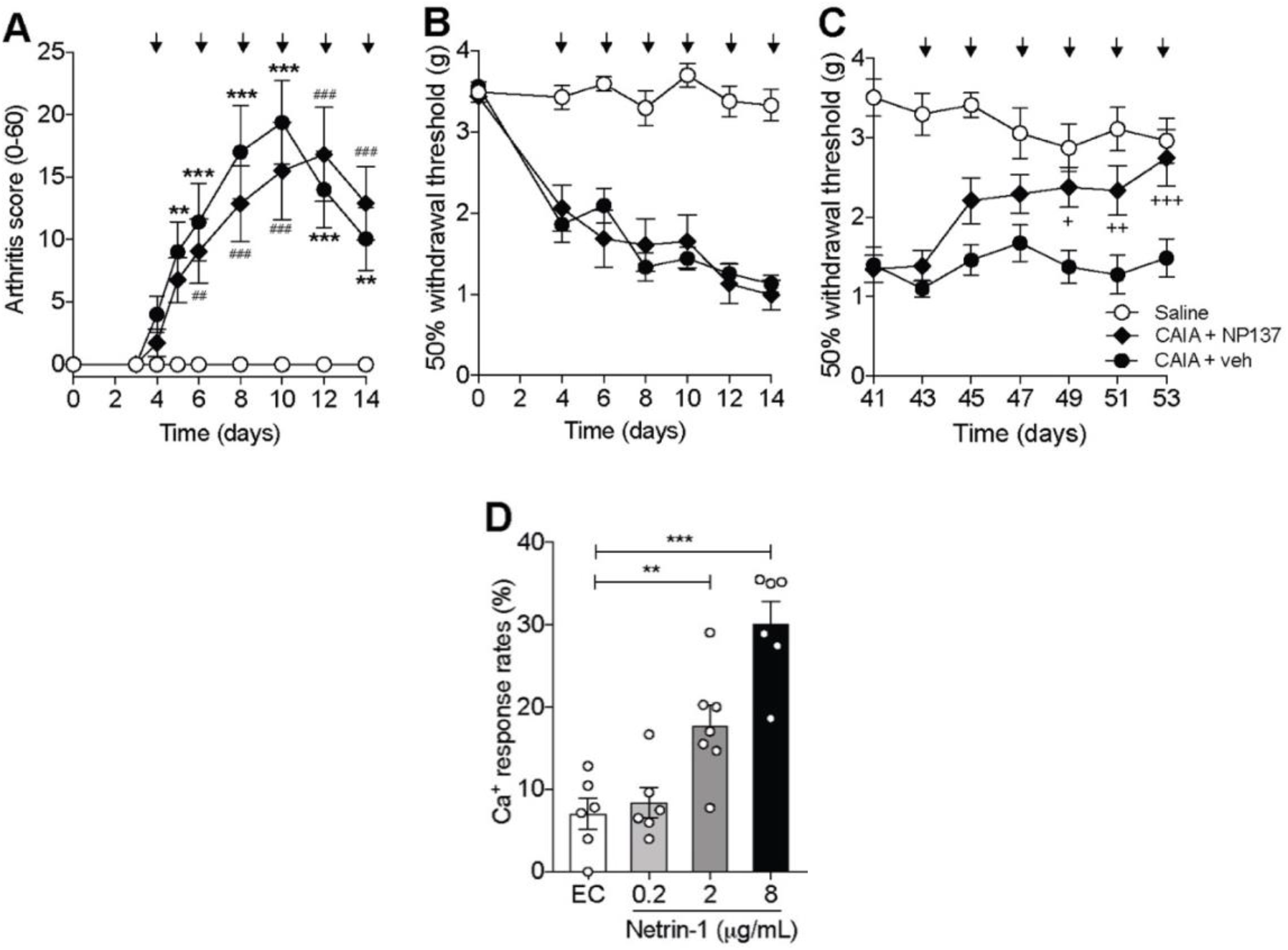
Netrin-1 has pronociceptive properties in the collagen antibody-induced arthritis (CAIA) model. Line graphs depict arthritis score of the hind ankle joints **(A)** and mechanical withdrawal thresholds **(B, C)** following intraperitoneal injection of the anti-netrin-1 antibody (NP137, 10 mg/kg) during the inflammatory (day 4-14) and late phase (day 43-53). Bar graphs represent intracellular [Ca^2+^] signal response rate in DRG neuronal cultures following stimulation with mouse recombinant netrin-1 protein (0.2-8 μg/mL) or extracellular solution (EC) as control **(D)**. Data are presented as mean ± SEM, n=9-10 mice/group **(A-C)** and n=6-7 technical replicates **(D)** ^*,+^p<0.05, ^**,##,++^p<0.01, ^***,###,+++^p<0.001. For **Figure 7A-C**, *represents saline vs. CAIA + vehicle, ^#^represents saline vs. CAIA + NP137 and ^+^represents CAIA + NP137 vs. CAIA + vehicle. Arrows indicate drug administration

## DISCUSSION

Here, we provide evidence that osteoclasts play important roles in CAIA-induced nociception, particularly in the refractive state. Strikingly, while osteoclast activity was predominantly enhanced during the inflammatory phase, pain-like behavior was reversed by osteoclast inhibitors, zoledronate and T06, only in the late phase. We found that both vascular and nerve densities were increased in the calcaneus during inflammation, which remained increased in the late phase despite resolution of joint inflammation. Importantly, the CAIA-induced refractive changes in bone, vascular and nerve densities in the late phase were attenuated by both osteoclasts inhibitors through reduced production of osteoclast-derived angiogenic and neurotrophic factors.

Osteoclast-induced acidification is frequently suggested as the primary mechanism linking osteoclasts to bone pain [10, 11, 30]. We present the possibility that osteoclasts contribute to pain via different mechanisms during periods of low bone resorptive activity, such as after resolution of joint inflammation in the CAIA model. It is unlikely that acidification is the only mechanism in which osteoclasts mediate pain. First, the resorption pit is tightly sealed by integrins to prevent destruction of nearby bone structures [9]. Thus, unless there is long-lasting sensitization of nociceptors that is only transiently mediated by protons, there must be a constant proton leakage from the resorption lacunae to be able to sensitize nociceptors in the bone. Second, reports show that osteoclast-blocking agents reverse pain-related behaviors in models without clear bone pathologies [12–14]. While one could argue that this is due to off-target effects, it also open up for the prospect that osteoclasts are secreting pronociceptive factors without resorbing bone. Emerging data show multifunctional roles of osteoclasts [15–19] and in support of this notion, we identified potential nociceptive roles of PDGF-BB, Sema4D and netrin-1 released by osteoclasts. While the exact mechanisms as to how these factors induce pain in the CAIA model remains unclear, our findings raise several possibilities by which osteoclasts may mediate nociception. For example, they could i) directly activate sensory neurons by release of netrin-1, ii) modulate neurite outgrowth/fasciculation, and/or iii) induce vascular growth, which in turn supports neoinnervation [31].

Interestingly, we found that it was not necessary to target all aspects of osteoclast activity to reduce pain as T06 only reduced PDGF-BB levels released from osteoclasts, whereas zoledronate also decreased Sema4D and netrin-1 levels. Furthermore, targeting netrin-1 alone was sufficient to produce similar antinociceptive effects as the osteoclast inhibitors in the late phase. Our work adds to the growing evidence that netrin-1 contributes to bone-associated pain [15, 28, 29] by directly exerting its action on nociceptors. Although the exact mechanism of action remains to be determined it is intriguing that mRNA for at least one of the two netrin-1 receptors, UNC5B, is detected in DRG neurons in two publicly available RNA-seq datasets [32, 33]. To our knowledge, however, our study is the first to report that blockade of netrin-1 led to reversal of pain-like behavior in an experimental model.

The temporal profile of bone destruction and pain-related behaviors during the inflammatory phase of the CAIA model coincides with clinical events observed in RA patients [34, 35]. Furthermore, we detected signs of bone erosions and persistent hypersensitivity in the late phase, which supports clinical data showing that residual signs of bone erosion are still detected in RA patients with good disease control [36, 37]. However, we are not aware of studies that correlates this to the remaining pain observed in subsets of patients. Moreover, we did not observe an anti-inflammatory effect of T06 as previously reported [38], which could be a model or treatment time-dependent difference. Nonetheless, our data show that two osteoclast inhibitors attenuated refractive hypersensitivity in the CAIA model without affecting joint inflammation. While there are no comparable clinical studies on RA-induced pain, reports show beneficial effects of osteoclast inhibition on other bone pain conditions disorders such as osteoarthritis, osteoporosis and bone metastatic cancer [6, 7, 39].

This study has some limitations. We did not evaluate other pain-related behaviors, such as heat hypersensitivity, which have been shown to be reversed by zoledronate in the glucose-6-phosphate isomerase-induced arthritis model [8]. Also, we acknowledge that zoledronate and T06 may have off-target effects as the pain-reliving properties of these drugs have been associated with actions in the CNS [12, 40]. Of importance, however, our data show that two drugs with different mechanisms induce bone protective effects [41–43] and reverse hypersensitivity, and thus antinociceptive effects are likely mediated via osteoclasts.

In conclusion, we provide novel insights to the pronociceptive actions of osteoclasts and suggest that they are not only linked to their resorptive activity but also coupled to their ability to modulate bone vascularization, bone innervation and release of factors such as netrin-1. Our findings therefore suggest a potential analgesic benefit of osteoclasts inhibition in RA patients with low disease activity. A netrin-1 antibody is currently clinically assessed in cancer indications [44, 45] and has demonstrated a favorable safety profile that could be compatible with investigating the clinical relevance of netrin-1 interference in RA-induced pain.

## Acknowledgements

We appreciate the technical assistance provided by Birgit McDonald and Kaja Sondergaard Laursen for *in situ* hybridization and immunohistochemistry. We would also like to thank the IVIA platform (UCA) for assistance in scintigraphic imaging.

## Contributors

RR and CIS designed the experiments, analyzed the data and wrote the manuscript along with input from all co-authors. RR performed behavior experiments, osteoclast assays and molecular tissue analyses. NA performed behavior experiments. ABF performed DRG culture experiments and calcium imaging. JB and FM performed CAIA experiments and scntigraphic imaging. AMM, EMI and JMJA conducted micro-CT imaging analyses. AB and TLA assisted in histomorphometric analyses. PP and DB provided cathepsin K inhibitor tanshinone IIA sulfonic sodium (T06) and PM provided anti-Netrin-1 antibody. All authors reviewed and approved the final version of the manuscript.

