## Supplementary information for "Bone innervation and vascularization regulated by osteoclasts contribute to refractive pain-related behavior in the collagen antibody-induced arthritis model"

***Drug delivery***

Zoledronate (100 μg/kg, Sigma-Aldrich) was injected subcutaneously (s.c.) every 3 days on day 6-56, 6-19 or 43-56 of the CAIA model. Phosphate-buffered saline (PBS) was used as vehicle control. The cathepsin K inhibitor tanshinone IIA sulfonic sodium (T06) was kindly provided by Dr. Dieter Brömme (University of British Columbia, Canada). T06 (40 mg/kg) was administered daily per oral gavage on day 6-21 or 43-59 of the CAIA model and filtered distilled water was used as vehicle. Anti-Netrin-1 antibody (Net1-mAb, NP137) was kindly provided by Netris Pharma (Lyon, France). NP137 (10 mg/kg) was injected i.p. every other day on day 4-14 or 43-51 of the CAIA model. Control buffer (containing 20 mM Tris, 5% sucrose and 0.01% Tween-20) was used as vehicle. All drug doses were chosen based on previous studies [1-3].

***Micro-computed tomography imaging***

Mice were euthanized with decapitation under deep isoflurane anesthesia (4%). Hind ankle joints were harvested, post-fixed in 4% paraformaldehyde (PFA) for 48 h, washed with 0.1 M PBS and stored at 4°C until further analysis. Distal tibia, talus and calcaneus were analyzed using a micro-computed tomography (microCT) imaging system (Skyscan 1272, Bruker). The scanning process was conducted at 10 µm voxel size, 60 kVp/166 µA X-Ray power and 627 ms integration time. Acquired images were reconstructed using NRecon software (Bruker) and analyzed using CT analyzer program (Bruker).

***^99m^Tc-HMDP scintigraphic imaging***

To evaluate bone remodelling, ^99m^Tc-HMDP scintigraphic imaging was performed on day 6, 17, 31 and 49 of the CAIA model. Mice were injected i.p. with ^99m^Tc-HMDP radiotracer (10 MBq/mouse, Osteocis®, IBA) and imaging was acquired 2.5 h post-injection. Planar acquisition was performed for 5 minutes (15% window at 140 keV) on the animal positioned in ventral decubitus on a parallel collimator (20mm/1.8/0.2) of a gamma camera for small animal (γImager®, Biospace) under isoflurane anaesthesia (1%). Quantitative analysis of bone scintigrams was performed with Gammavision*+* software (Biospace) using fixed-sized rectangular regions of interest on both hind paws. For each time point and animal, scintigraphic ratio of bone (SR_b_) were both calculated as follow: SR = pathological paw average activity (cpm)/contralateral paw average activity (cpm).

***Immunoassay***

Mice were deeply anesthetized with isoflurane (4%) and blood was collected with cardiac puncture. Blood was then allowed to clot for 30 min at room temperature, centrifuged (4800 *g* for 10 min at 4^o^C) and the collected serum stored at -80^o^C until analyses. Serum levels of C-telopeptide of type I collagen (CTX-I, bone erosion marker) and propeptide of type I procollagen (PINP, boner formation marker) were quantified in mouse serum using the RatLaps CTX-I EIA and Rat/Mouse PINP EIA kits (IDS). Samples and standards were measured in duplicates.

***Tissue processing for paraffin-embedded sections***

Animals were deeply anesthetized with isoflurane (4%) and transcardially perfused with PBS followed by 4% PFA. Hind ankle joints were harvested and postfixed for 48 h in 4% PFA. Next, tissues were washed in PBS and decalcified in 10% ethylenediaminetetraacetate (EDTA) solution (Sigma-Aldrich) for 4-5 weeks. Decalcification solution was changed every week. The decalcified joints were then dehydrated using 70% ethanol and xylene followed by embedding in paraffin. Sagittal sections (5 μm) were cut using a rotary microtome (Microm HM360, Thermo Fisher) and mounted onto glass slides.

***Immunohistochemistry***

Prior to staining, paraffin-embedded sections were deparaffinized, blocked for endogenous peroxidase and subjected to overnight antigen retrieval in Tris-acetate buffer (pH 6, DAKO) at 60^o^C. The following day, sections were blocked with 5% normal goat serum to reduce nonspecific binding and incubated with primary antibodies (cathepsin K, 1:100, OriGene TA318059; PGP9.5, 1:500, Cedarlane 184; CD31, 1:250, Abcam ab124432) for 2 h. Subsequently, alkaline phosphatase-conjugated IgG (1:500, Sigma-Aldrich) was used for secondary detection followed by visualization using liquid fast-red substrate kit (Abcam). All sections were counterstained with Mayer’s hematoxylin and mounted with ProlongGold antifade mountant (ThermoFisher).

***In situ hybridization***

Paraffin-embedded sections were in situ hybridized using a modified version of the RNAScope 2.0 high-definition procedure (ACD) [4] to detect mRNA expression of *Acp5* using 20 double Z probe pairs targeting nucleotide 200-1414 of *Acp5* ([NM_001102405.1](http://www.ncbi.nlm.nih.gov/nuccore/NM_001102405.1)). In short, sections were incubated with pretreatment 1 for 10 min at room temperature, pretreatment 2 for 15 min at 85^o^C and pretreatment 3 was replaced with a pepsin treatment (10% diluted in RNase-free water, DAKO) for 20 min at 40^o^C. The subsequent hybridization and amplification steps were performed according to the manufacturer’s instructions. Next, sections were incubated with digoxigenin (DIG)-labeled tyramide (1:200) from TSA plus DIG reagent kit (PerkinElmer) for 5 min at room temperature followed by labelling with alkaline phosphatase-conjugated anti-DIG FAB fragments (1:1000, Roche). Labelled sections were visualized with liquid permanent red (DAKO), counterstained with Mayer’s hematoxylin and mounted with Aquatex (Sigma-Aldrich).

***Histomorphometry***

Sections were scanned using a NanoZoomer Slide Scanner (Hamamatsu Photonics) and images were analyzed using NDP.view2 software (Hamamatsu Photonics). The conventional histomorphometric parameter osteoclast surface per bone surface (Oc.S/BS) was estimated using a grid of sinus curves. CTSK- and *Acp5*-stained cells were quantified at intersection points between sinus curves and bone surfaces. Bone vascular and nerve densities were estimated by quantifying CD31- and PGP9.5-stained vessels or nerves in areas located below 100 μm from bone surfaces. A point-grid was used to estimate the size of the tissues and local vessel and nerve densities were expressed as the number of vessels or nerves per tissue area (mm^2^). Analysis was performed by investigators who were blinded to the group assignments.

***Quantitative real-time polymerase chain reaction***

Mice were euthanized with decapitation under deep isoflurane anesthesia (4%). Hind ankle joints and L3-L5 dorsal root ganglia (DRGs) were harvested, flash-frozen and stored at -80^o^C until further analysis. mRNA from these tissues was extracted using TRIzol reagent (Invitrogen) according to the manufacturer’s protocol followed by cDNA synthesis using MultiScribe Reverse Transcriptase (Invitrogen). qPCR was performed using StepOne Real-Time PCR Systems (Applied Biosystems) using pre-developed Taqman primer/probe sets: *Acp5* (Mm00475698_m1), *Asic3* (Mm00805460_m1), *Clcn7* (Mm00442400_m1), *Ctsk* (Mm00484039_m1), *Ntn1* (Mm00500896_m1), *Pecam1* (Mm01242576_m1), *Sema4d* (Mm00443147_m1), *Tcirg1* (Mm00469406_m1), *Trpv1* (Mm01246402_m1), *Hprt1* (Mm03024075_m1) and *Rplp2* (Mm00782638_m1). Threshold cycle values for each sample were used to calculate the number of cell equivalents in the test samples using the standard curve method. The data were normalized to mRNA levels of housekeeping genes (*Hprt1* or *Rplp2*) and expressed as relative expression.

***Bone marrow-derived primary osteoclast culture***

Bone marrow was flushed from mouse tibia and femur using 27-gauge needle, and cells were isolated by centrifugation at 300*g* for 5 min. Cells were then cultured in Eagle’s Minimum Essential Medium - alpha modification (αMEM, Gibco) supplemented with 10% fetal bovine serum (FBS, Sigma-Aldrich), 1% penicillin/streptomycin (ThermoFisher), 30 ng/mL macrophage colony-stimulating factor (M-CSF, R&D) and 50 ng/mL receptor activator of nuclear kappa-B ligand (RANKL, R&D) at a density of 1.25 x 10^5^ cells/well for 6 days in a 37°C incubator with 5% CO_2._ Cells cultured without RANKL, which prevents osteoclast differentiation and would instead yield monocytes/macrophages, were included as control. Medium was refreshed every 2 days and starting on day 2 cells were stimulated with either 600 nM zoledronate (Sigma-Aldrich), 1 µM T06 or vehicle (PBS). On the day of medium change, culture supernatant was collected and stored at -80^o^C for further analysis. At the end of experiment, cells were fixed in 4% PFA for 5 min and stained for tartrate resistant acid phosphatase (TRAP) using leukocyte acid phosphatase kit (Sigma-Aldrich). In short, fixed cells were incubated with the following reagents: Fast Garnet GBC, sodium nitrite, naphthol AS-BI phosphoric acid, tartrate and acetate solution for 30-45 min at 37^o^C. Quantification of multinucleated cells (>3 nuclei) was performed in three independent experiments using a brightfield microscope (Nikon Eclipse TE-200).

***Western Blot***

Netrin-1 protein levels were assessed by Western blotting of whole osteoclast lysate harvested on day 6 since this protein is an extracellular matrix protein and therefore does not diffuse into solution [2]. Osteoclasts were homogenized in lysis buffer containing PBS, 1% sodium dodecyl sulphate (SDS) and a cocktail of protease inhibitors. 5 μg of proteins were separated using gel electrophoresis and transferred onto nitrocellulose membranes. Nonspecific binding sites were blocked with 5% non-fat milk, and the membranes was probed with primary antibody overnight at 4^o^C (netrin-1, 1:1000, Abcam ab1267729). After washing, the membranes were incubated with secondary antibodies conjugated to HRP (CST) and chemiluminescent reagent (SuperSignal West Femto, ThermoFisher) was used to visualize immunopositive band on ChemiDoc Imaging System (Biorad). Membranes were then stripped and reprobed with primary antibody as protein reference (β-actin, 1:10000, CST 3700). Quantification was performed with ImageJ, and results are presented as percentage change to the control group.

***Calcium imaging***

Cells were loaded with the calcium indicator Fluo-3 AM (4.4 μM, ThermoFisher) for 30 min at room temperature and washed with extracellular solution (EC; 10 mM Hepes, 2 mM CaCl_2_, 3 mM KCl, 145 mM NaCl, 2 mM MgCl_2_, 10 mM glucose, pH 7.4). Calcium imaging was then performed with Zeiss LSM800 confocal microscope with the following settings: 20x objective, argon laser (488 nm excitation), change in emission (506 nm) measured every 2.5 sec using a photomultiplier tube. Mouse recombinant netrin-1 (0.2, 2 or 8 μg/mL, R&D) was directly applied to the cells and cells responses were recorded for 16 min. EC solution was used as negative control for the experiment. At the end of each experiment, KCl (50 mM) was applied to detect functional neurons. Acquired images were analyzed using ImageJ. In each image, the mean fluorescence intensity (F) of each neuron was measured manually selecting all cells present in the field of image. The baseline recording (F_0_) of each neuron was calculated as the average mean signal of the initial 10 images of the series before application of reagents. A cell was considered positive if the increase of the fluorescent signal was at least ≥25% compared to baseline. Data are presented as response rates for each dish (positive cells/total cells).

**SUPPLEMENTARY TABLES AND FIGURES**

**Table S1.** Bone parameters evaluated by micro-CT imaging in distal tibia and talus of saline- and CAIA-subjected mice

|  | **Inflammatory** | | | **Late** | | |
| --- | --- | --- | --- | --- | --- | --- |
|  | **Saline (n=6)** | **CAIA (n=5)** | **p-value** | **Saline (n=6)** | **CAIA (n=6)** | **p-value** |
| **Distal tibia** |  | | | | | |
| tBMD (g/cm^3^) | 3.957 ± 0.604 | 3.343 ± 1.419 | 0.358 | 0.037 ± 0.007 | 0.046 ± 0.003 | 0.008** |
| BV/TV (%) | 0.543 ± 0.522 | 0.115 ± 0.116 | 0.108 | 0.376 ± 0.227 | 0.342 ± 0.447 | 0.875 |
| Tb.Th (mm) | 0.050 ± 0.024 | 0.037 ± 0.016 | 0.342 | 0.049 ± 0.026 | 0.035 ± 0.027 | 0.383 |
| Tb.N (1/mm) | 0.085 ± 0.061 | 0.025 ± 0.018 | 0.062 | 0.053 ± 0.040 | 0.029 ± 0.050 | 0.395 |
| Tb.Sp (mm) | 0.410 ± 0.025 | 0.441 ± 0.026 | 0.078 | 0.430 ± 0.044 | 0.423 ± 0.028 | 0.759 |
| cBMD (g/cm^3^) | 0.388 ± 0.082 | 0.318 ± 0.152 | 0.354 | 1.503 ± 0.019 | 1.511 ± 0.026 | 0.583 |
| Ct.Th (mm) | 0.312 ± 0.015 | 0.300 ± 0.014 | 0.130 | 0.300 ± 0.015 | 0.307 ± 0.019 | 0.499 |
| Ct.Ar (mm) | 0.692 ± 0.050 | 0.724 ± 0.001 | 0.193 | 0.675 ± 0.050 | 0.737 ± 0.024 | 0.021* |
| **Talus** |  | | | | | |
| tBMD (g/cm^3^) | 0.388 ± 0.008 | 0.318 ± 0.152 | 0.354 | 0.371 ± 0.094 | 0.247 ± 0.104 | 0.069 |
| BV/TV (%) | 28.62 ± 6.276 | 22.21 ± 13.10 | 0.129 | 27.04 ± 7.234 | 17.87 ± 7.784 | 0.074 |
| Tb.Th (mm) | 0.072 ± 0.005 | 0.061 ± 0.018 | 0.189 | 0.068 ± 0.007 | 0.068 ± 0.009 | 0.939 |
| Tb.N (1/mm) | 3.957 ± 0.604 | 3.343 ± 1.419 | 0.358 | 3.935 ± 0.721 | 2.554 ± 0.961 | 0.018* |
| Tb.Sp (mm) | 0.211 ± 0.022 | 0.229 ± 0.087 | 0.617 | 0.202 ± 0.030 | 0.265 ± 0.078 | 0.102 |

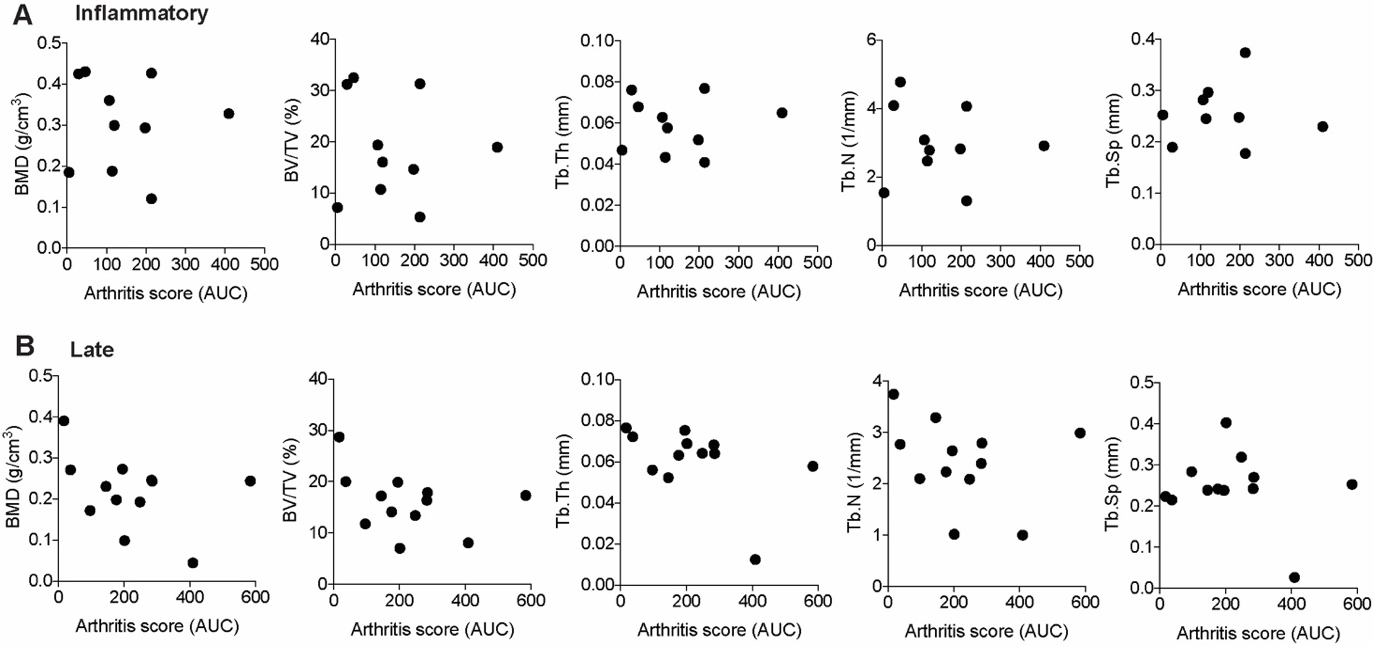

**Figure S1**. Arthritis score does not correlate with the degree of bone erosion in CAIA animals. Scatter graphs depict the relationship between arthritis scores (presented as area under the curve, AUC) vs. bone mineral density (BMD), bone volume per tissue volume (BV/TV), trabecular thickness (Tb.Th), trabecular number (Tb.N) and trabecular separation (Tb.Sp) of calcaneus from inflammatory (**A**) and late (**B**) phase CAIA-subjected mice. AUC was calculated from arthritis scores recorded throughout the inflammatory phase of each mouse
